## Supplementary figures and images for "Divergent selection following speciation in two ectoparasitic honey bee mites"

### Supplementary Figures 1 to 7

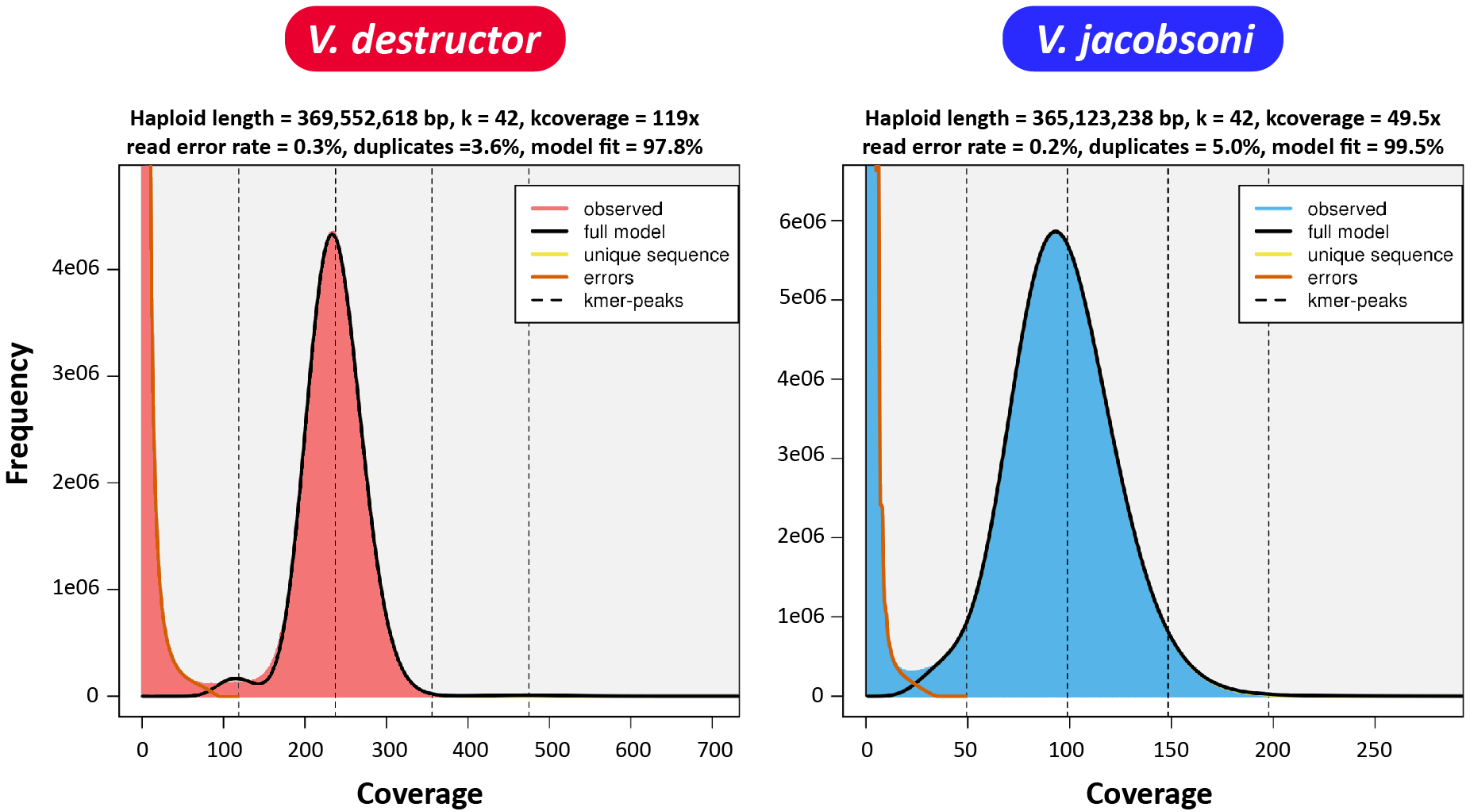

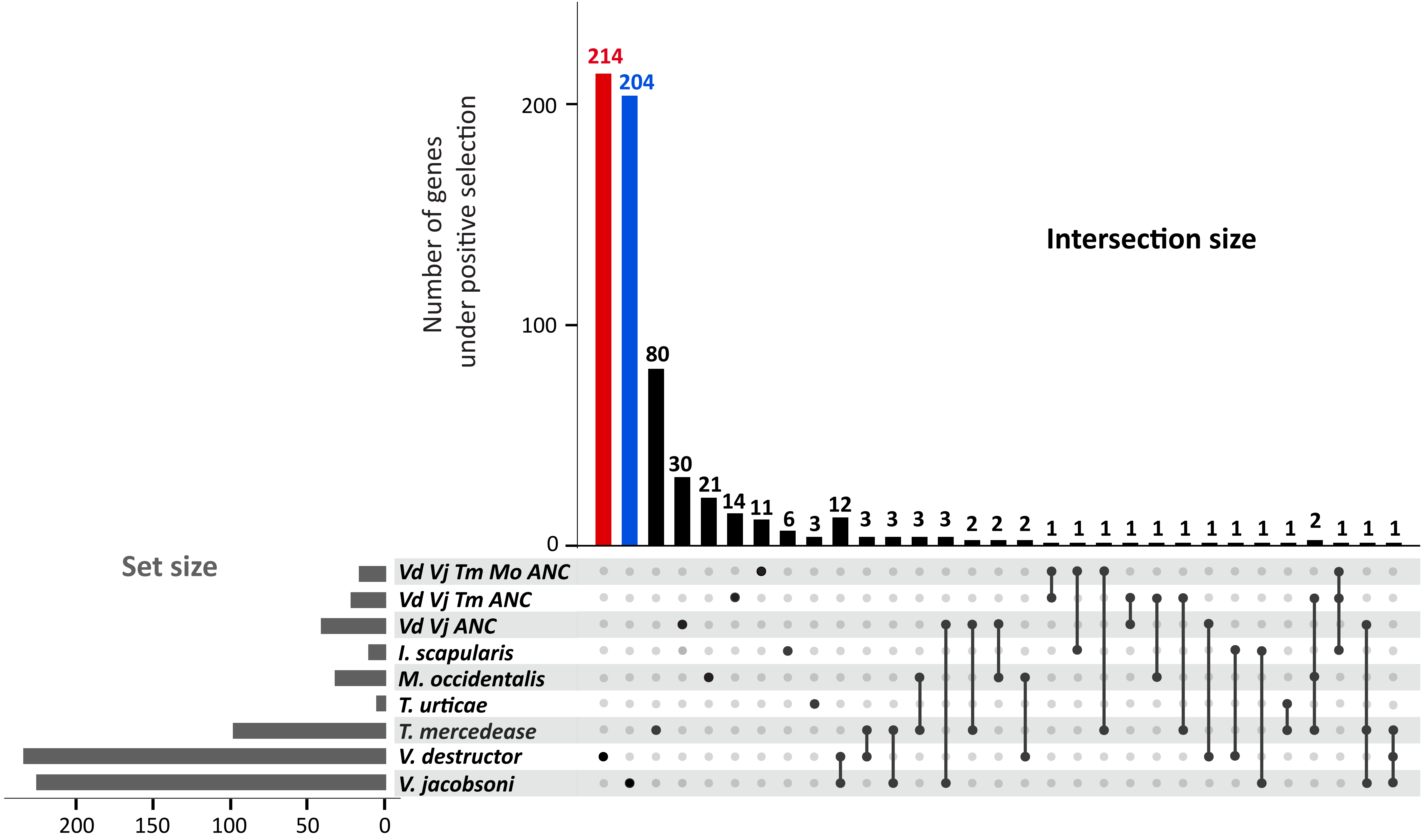

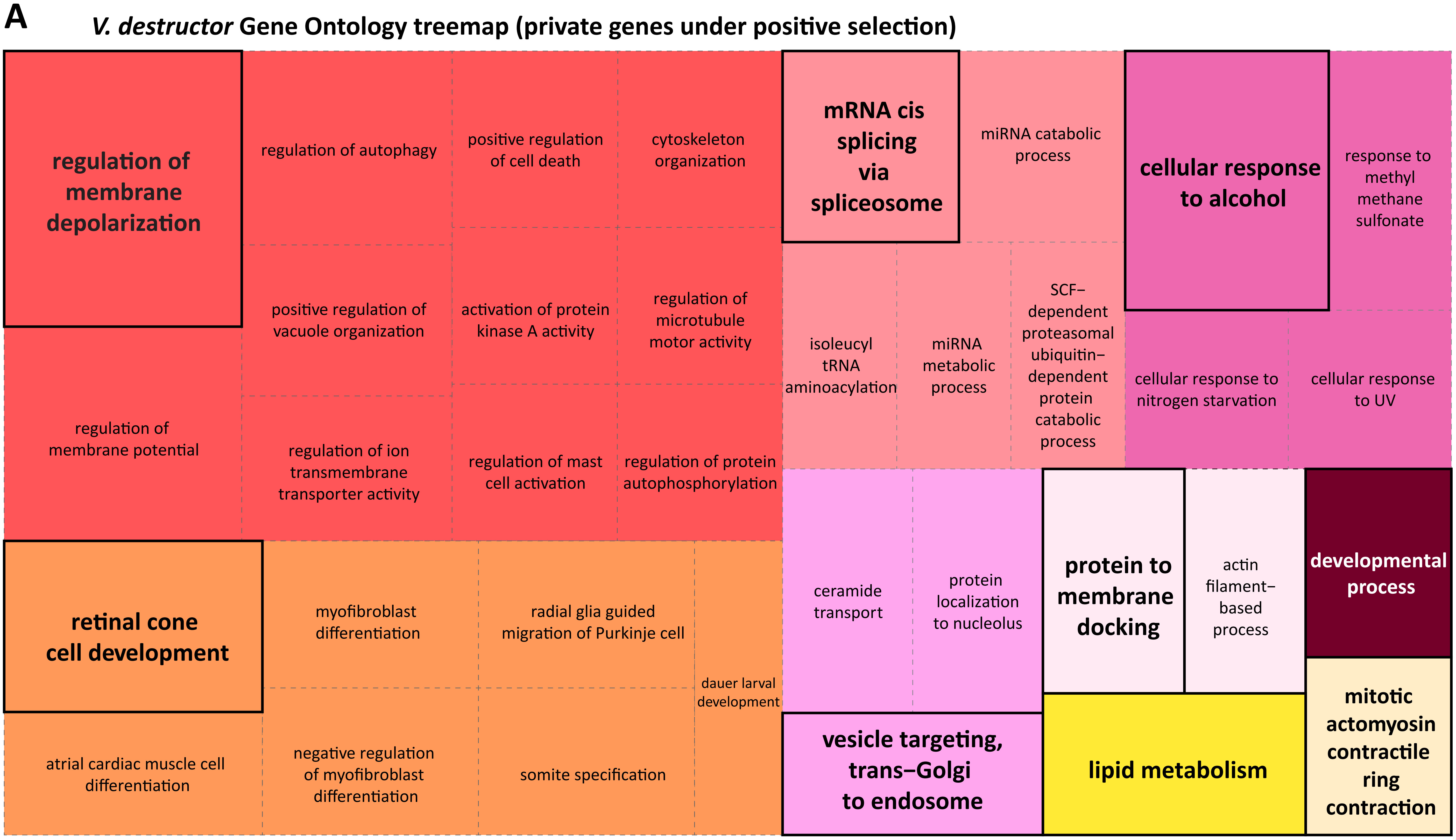

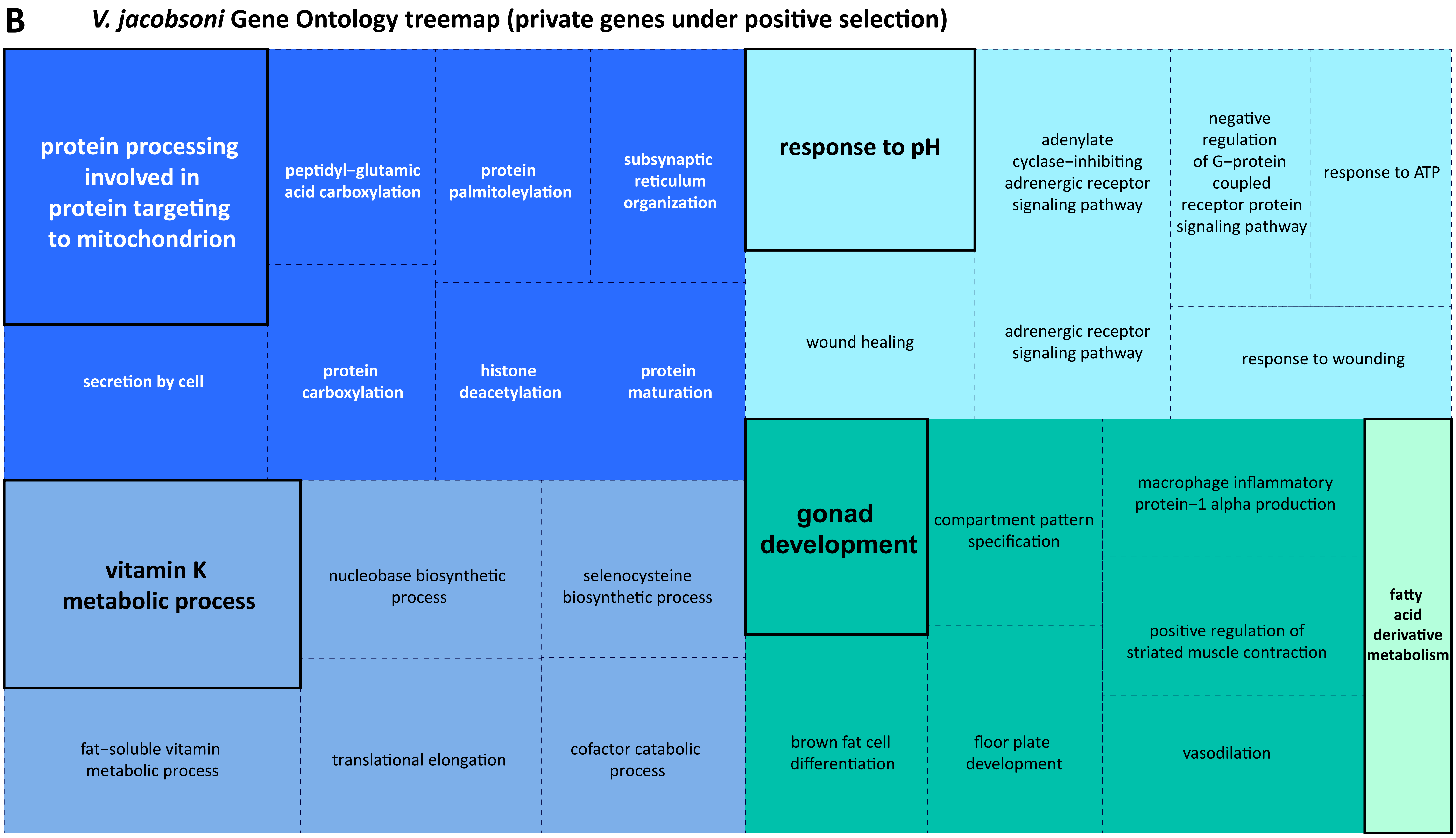

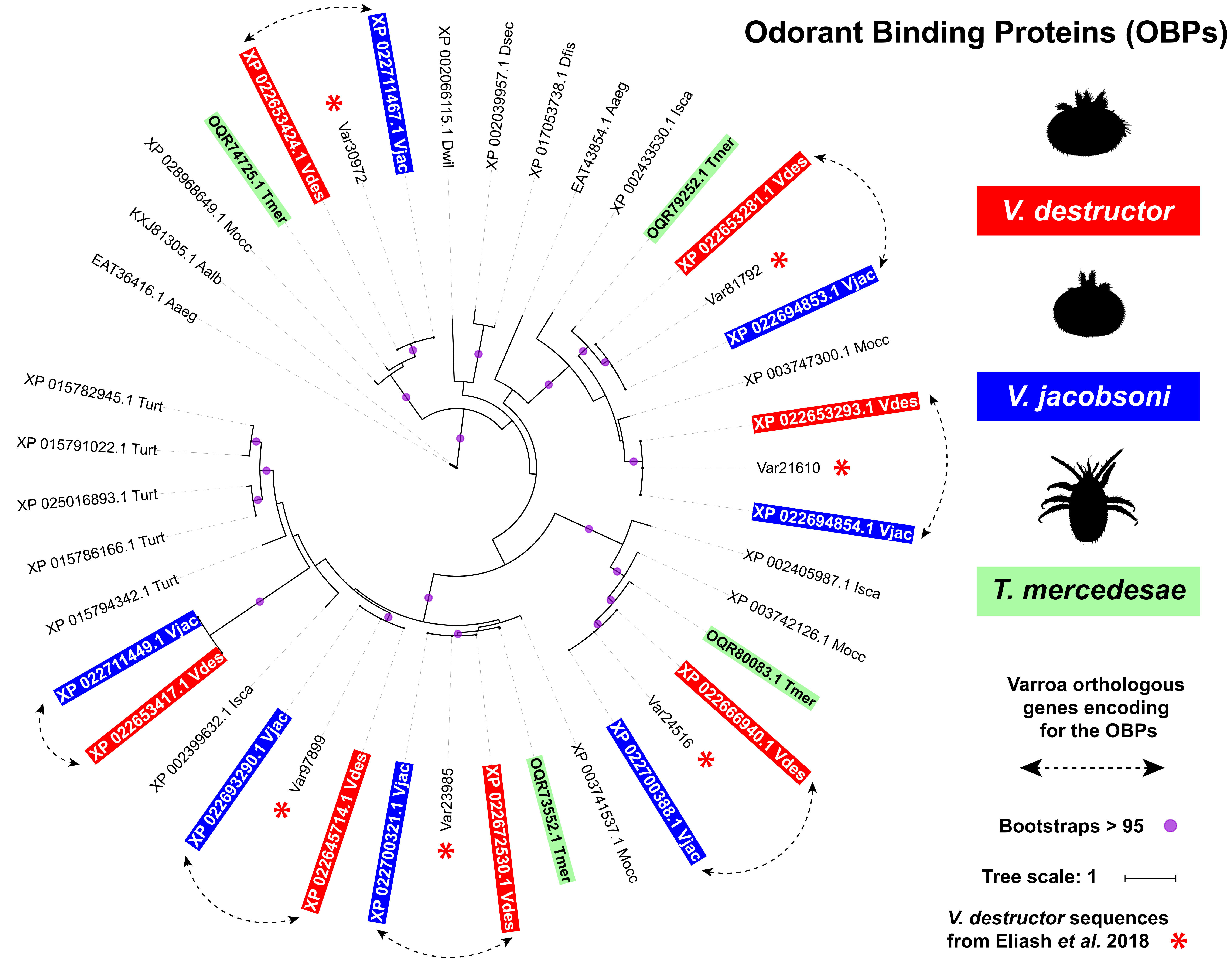

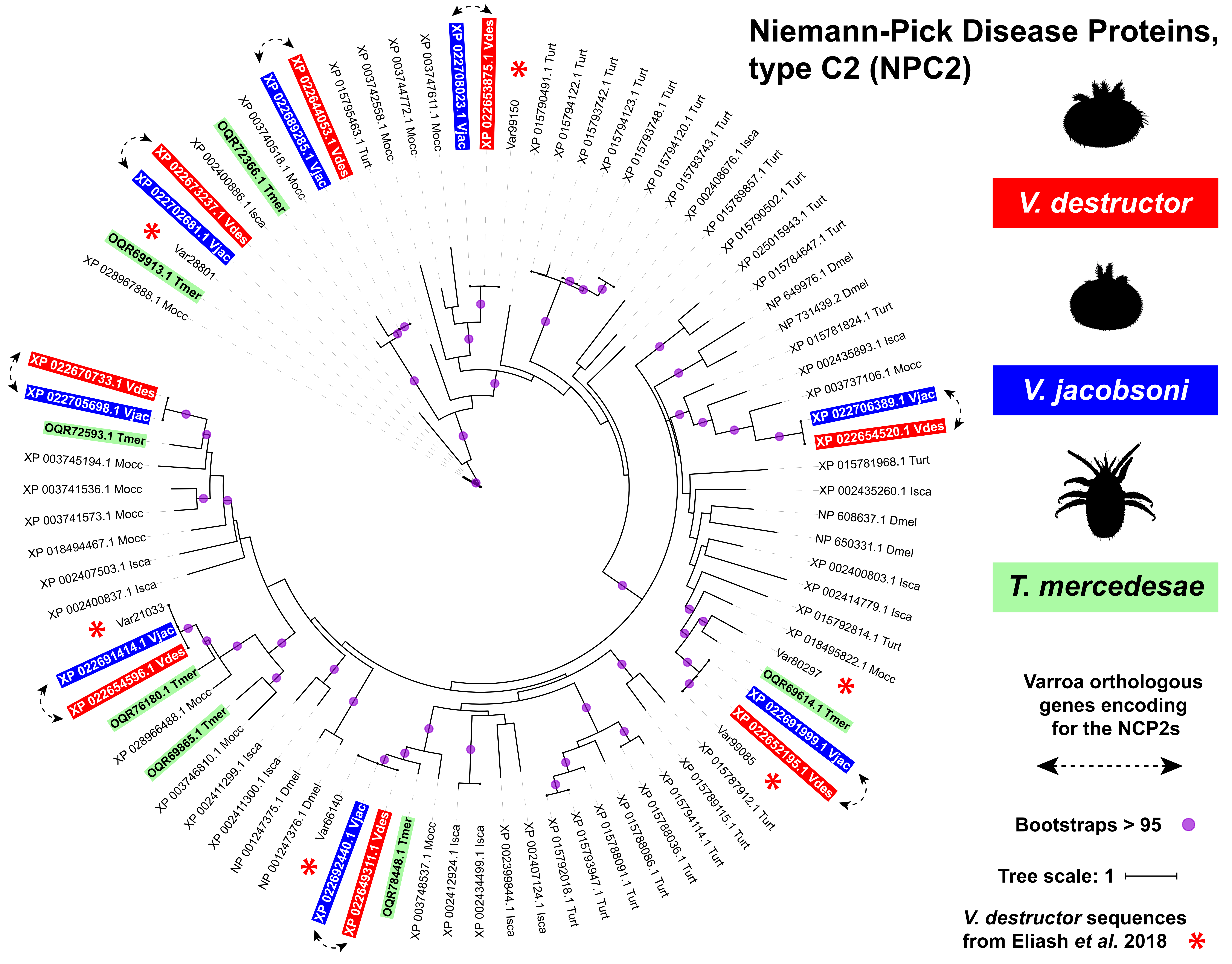

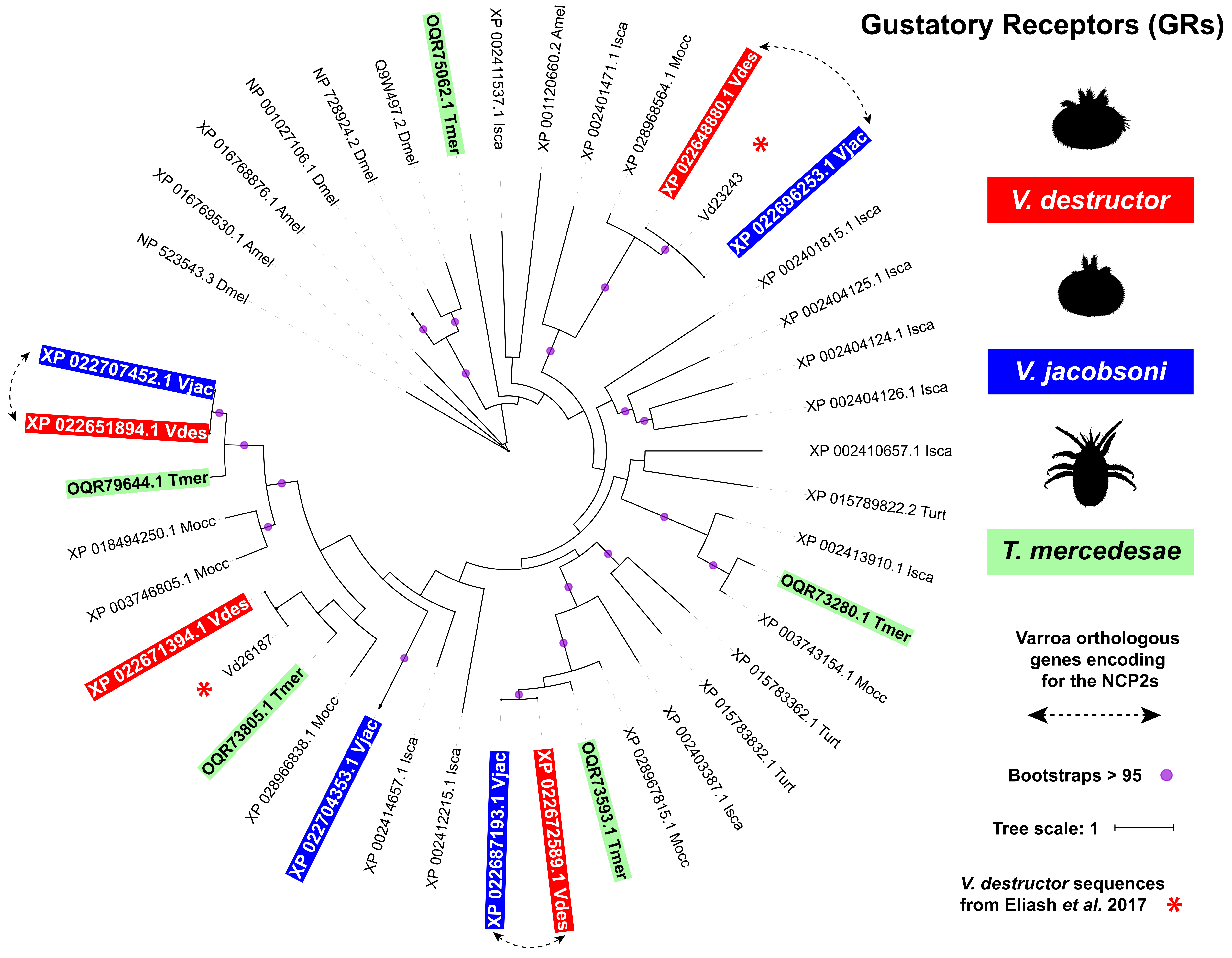

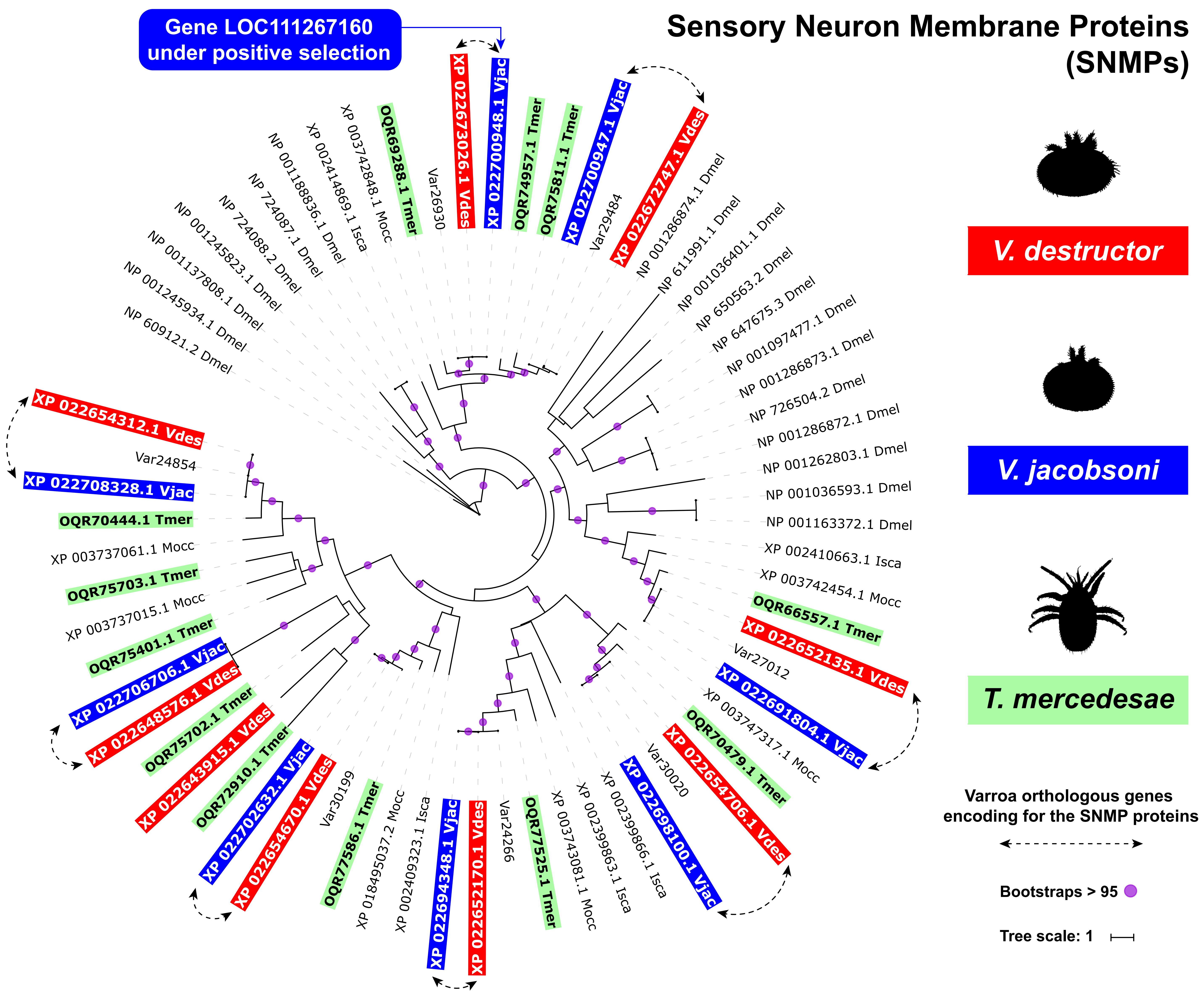
